# The Lack of Nrf2 Causes Hepatocyte Dedifferentiation and Reduced Albumin Production in an Experimental Extrahepatic Cholestasis Model

**DOI:** 10.1101/2021.04.26.441515

**Authors:** Guo-Ying Wang, Veronica Garcia, Joonyong Lee, Jennifer Yanum, Huaizhou Jiang, Guoli Dai

## Abstract

The transcription factor Nrf2 modulates the initiation and progression of a number of diseases including liver disorders. The aim of this study was to evaluate whether Nrf2 mediates hepatic adaptive responses to cholestasis. Wild-type and Nrf2-null mice were subjected to bile duct ligation (BDL) or a sham operation. Various assessments were performed at different days after surgery. Significant genotype-dependent changes in liver size, biliary ductular reaction, hepatocyte proliferation, and fibrotic response were not observed. However, as cholestasis progressed to Day 15 post-BDL, hepatocytes in the wild-type mice exhibited a tendency to dedifferentiate, indicated by the very weak expression of hepatic progenitor markers: CD133 and fibroblast growth factor-inducible 14 (Fn14). During the same period, Nrf2 deficiency augmented this tendency, manifested by higher CD133 expression, earlier, stronger, and continuous induction of Fn14 expression, and markedly reduced albumin production. Remarkably, as cholestasis advanced to the late stage (40 days after BDL), hepatocytes in the wild-type mice exhibited a Fn14+ phenotype and strikingly upregulated the expression of deleted in malignant brain tumor 1 (DMBT1), a protein essential for epithelial differentiation during development. In contrast, at this stage, hepatocytes in the Nrf2-null mice entirely inhibited the upregulation of DMBT1 expression, displayed a strong CD133+/Fn14+ phenotype indicative of severe dedifferentiation, and persistently reduced albumin production. Collectively, our studies revealed that Nrf2 maintains hepatocytes in the differentiated state potentially via the increased activity of the Nrf2/DMBT1 pathway during cholestasis. These findings enable us to gain novel insight into how hepatocytes respond to cholestasis.

**New and Noteworthy:** We found that, when hepatocytes are exposed to cholestasis, they exhibit a tendency of dedifferentiation. In this case, Nrf2 is highly activated to markedly up-regulate the expression of epithelial differentiation gene DMBT1, which potentially prevent hepatocytes from dedifferentiation. Our findings revealed a plastic property of hepatocytes in response to cholestasis and demonstrated a novel Nrf2/DMBT1 pathway likely controlling this property of hepatocytes.

## Introduction

Nuclear factor erythroid 2-related factor 2 (Nrf2) is a leucine zipper motif-containing transcription factor ^1^. It primarily functions as a redox sensor and maintains redox homeostasis in a variety of organ systems. Upon activation, it exerts many beneficial effects, such as anti-oxidation, detoxification, anti-inflammation, and anti-apoptosis. During the past several decades, significant insights have been gained regarding its epigenetic and non-epigenetic regulation, activation mechanisms, target genes, and roles in health and disease ^2-5^. Nrf2 participates in modulating the initiation and progression of many pathological processes, including liver disorders, and has proven to be a promising target for the prevention and treatment of several diseases ^6^.

Obstructive extrahepatic cholestasis in humans is widely modeled in rodents by common bile duct ligation (BDL). BDL induces accumulation of the bile constituents, including bile acids, cholesterol, and bilirubin in the liver and blood, thereby causing oxidative stress and secondary liver injury ^7-9^. Nrf2 regulates the expression of genes that are critical not only for cellular defense against oxidative stress but also for bile acid synthesis and the enterohepatic circulation of bile acids, playing essential roles in maintaining bile acid homeostasis ^10^. During cholestasis, the liver exhibits adaptive responses, including Nrf2 activation, to alleviate liver injury ^10-12^. Genetic overactivation of Nrf2 via knockdown of its inhibitor Kelch-like ECH-associated protein 1 (Keap1) or pharmacological activation of Nrf2 counteracts the effects of cholestatic liver injury ^12,13^. However, several studies have demonstrated that the lack of Nrf2 does not cause worsened liver injury in BDL-induced extrahepatic cholestasis or alpha-naphthylisothiocyanate-induced intrahepatic cholestasis ^10-12^. These observations have contributed to Nrf2 absence-induced reductions in bile acid synthesis, excretion, and reabsorption, as well as an increase in bile acid hydroxylation and other unknown Nrf2-dependent mechanisms. Here, we aimed to gain further insights into how Nrf2 modulates hepatic adaptive responses to cholestasis.

## Materials and Methods

### Mice and surgical procedures

Nrf2+/+ and Nrf2-/-male mice (6-month-old) with a C57BL6/129SV mixed background were used ^1^. The mice were housed in plastic cages at 22 ± 1 °C on a 12 h light/dark cycle with lights on from 6:00 am to 6:00 pm. Standard rodent chow and water were provided *ad libitum* throughout the acclimatization period. The BDL and sham operation surgical procedures were performed under aseptic conditions. Mice were anesthetized using an isoflurane inhalation agent. The common bile duct was ligated with two ligatures separated by 2 mm. A cut was made between the two ligatures. Control mice underwent a sham operation that consisted of exposure but no ligation of the common bile duct. Mice were euthanized at 5, 10, 15, 25, and 40 days after the surgeries to collect their livers. The livers were immediately excised and weighed. A portion of each liver was fixed in 10% formalin and embedded in paraffin to prepare the liver sections. Meanwhile, part of each liver was frozen in liquid nitrogen to prepare the liver lysates. The remainder of each liver was frozen in liquid nitrogen and stored at -80 °C until further use. All animal experiments were conducted in accordance with the National Institutes of Health (NIH) Guide for the Care and Use of Laboratory Animals. Protocols for the care and use of animals were approved by the Indiana University-Purdue University Indianapolis Animal Care and Use Committee.

### Histology and immunohistochemistry

Formalin-fixed and paraffin-embedded liver sections were subjected to immunostaining. Primary antibodies against Ki-67 (BS1454; Bioworld, Irving, TX, USA), laminin (L-9393; Sigma-Aldrich, St. Louis, MO, USA), CD133 (14-1331; eBioscience, San Diego, CA, USA), CK19 (SC-33111; Santa Cruz Biotechnology, Dallas, TX, USA); NAD(P)H quinone dehydrogenase 1 (NQO1; A19586; ABclonal, Woburn, MA, USA), fibroblast growth factor-inducible 14 (Fn14; ab109365; Abcam, Cambridge, UK), and DMBT1 (AF5195; R&D Systems, Minneapolis MN, USA) were used. Ki67-postive hepatocytes were counted in five randomly chosen microscope fields per section at 200x magnification. Sirius Red-stained areas were quantified using Image-Pro Plus software (Media Cybernetics, Rockville, MD, USA).

### Western blot analysis

Liver homogenates (10 µg) were separated using polyacrylamide gel electrophoresis under reducing conditions. Proteins from the gels were electrophoretically transferred to polyvinylidene difluoride membranes. Antibodies against NQO1 (2618-1; Epitomics, Burlingame, CA, USA), CD133 (PAB12663; Abnova, Taipei, Taiwan), Fn14 (ab109365; Abcam), DMBT1 (AF5195; R&D Systems), phosphorylated yes-associated protein 1 (p-YAP; S127; 4911; Cell Signaling Technology, Danvers, MA, USA), albumin (A0353, Abclonal, Woburn, MA), YAP (14074; Cell Signaling Technology), p-mammalian target of rapamycin (p-mTOR; S2448; 4911; Cell Signaling Technology), mTOR (2983; Cell Signaling Technology), p-epidermal growth factor receptor (p-EGFR; Y1086; 1139-1; Epitomics), EGFR (1114; Epitomics), and β-catenin (9587; Cell Signaling Technology) were used as probes. Immune complexes were detected using an enhanced chemiluminescence system (Pierce, Rockford, IL, USA).

### Statistical analysis

Data are shown as mean ± standard deviation (SD). Statistical analysis was performed using one-way analysis of variance (ANOVA) or unpaired Student’s t-test. Significant differences were defined when *P* < 0.05.

## Results

### Nrf2 deficiency does not cause broadly worsened liver injury after BDL

We subjected wild-type and Nrf2-null mice to a sham operation or BDL and subsequently performed various assessments at different time points after the surgeries. We found that both genotype groups of mice exhibited markedly increased liver-to-body weight ratios at days 15 and 25 following BDL relative to their sham controls, respectively (Fig. 1). The data indicated that, regardless of Nrf2, the livers displayed dramatic enlargement, which was a robust response to BDL during this period (2-4 weeks post-BDL). Immunostaining for CK19, a marker of cholangiocytes, revealed similar biliary ductal responses to BDL between the two genotype groups of mice (Fig. 2). These findings suggested that Nrf2 might not play a crucial role in the adaptive response of the biliary ducts to cholestasis. Quantification of Ki-67^+^ hepatocytes showed that, irrespective of Nrf2, the hepatocytes underwent massive proliferation at day 5 after BDL, but this event was dramatically weakened thereafter (Fig. 3A and 3B), whereas a few replicating hepatocytes were seen in the sham controls (data not shown). Proliferating non-parenchymal liver cells were similarly concentrated in the biliary ductular areas of the wild-type and Nrf2-null mice subjected to BDL (Fig. 3A). These observations suggested that the presence or absence of Nrf2 might not affect the BDL-induced hepatocyte replication, the first line of repair response to liver injury. Laminin immunostaining showed hepatic septa formation, which was more evident after day 10 post-BDL, without an obvious Nrf2-dependent difference (Fig. 4A). Sirius Red staining reflected the hepatic content of collagen (Fig. 4B), which was equivalent between the BDL wild-type and Nrf2-null mice over the 40 day period (Fig. 4C). Taken together, these findings demonstrated that the lack of Nrf2 does not remarkably affect BDL-induced liver size alteration, biliary ductular reaction, liver repair, and hepatic fibrotic response.

**Figure 1.**
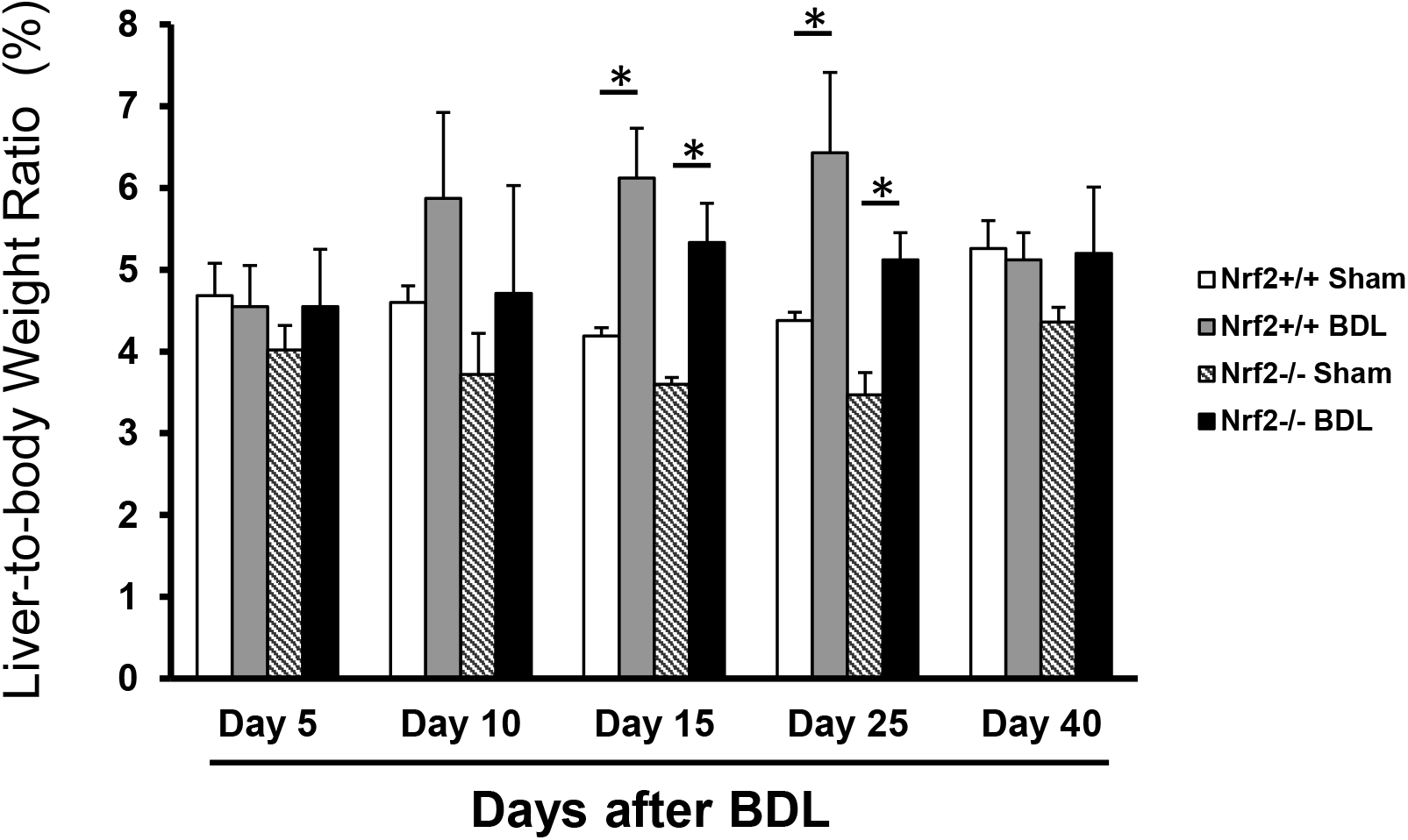
Liver-to-body weight ratios in Nrf2+/+ and Nrf2-/- mice after bile duct ligation (BDL) or sham operation (Sham). Adult male mice of both genotypes were subjected to BDL or Sham operation. Mice were sacrificed at the time points indicated. The liver-to-body weight ratios are shown. \**P* < 0.05 and n = 5.

**Figure 2.**
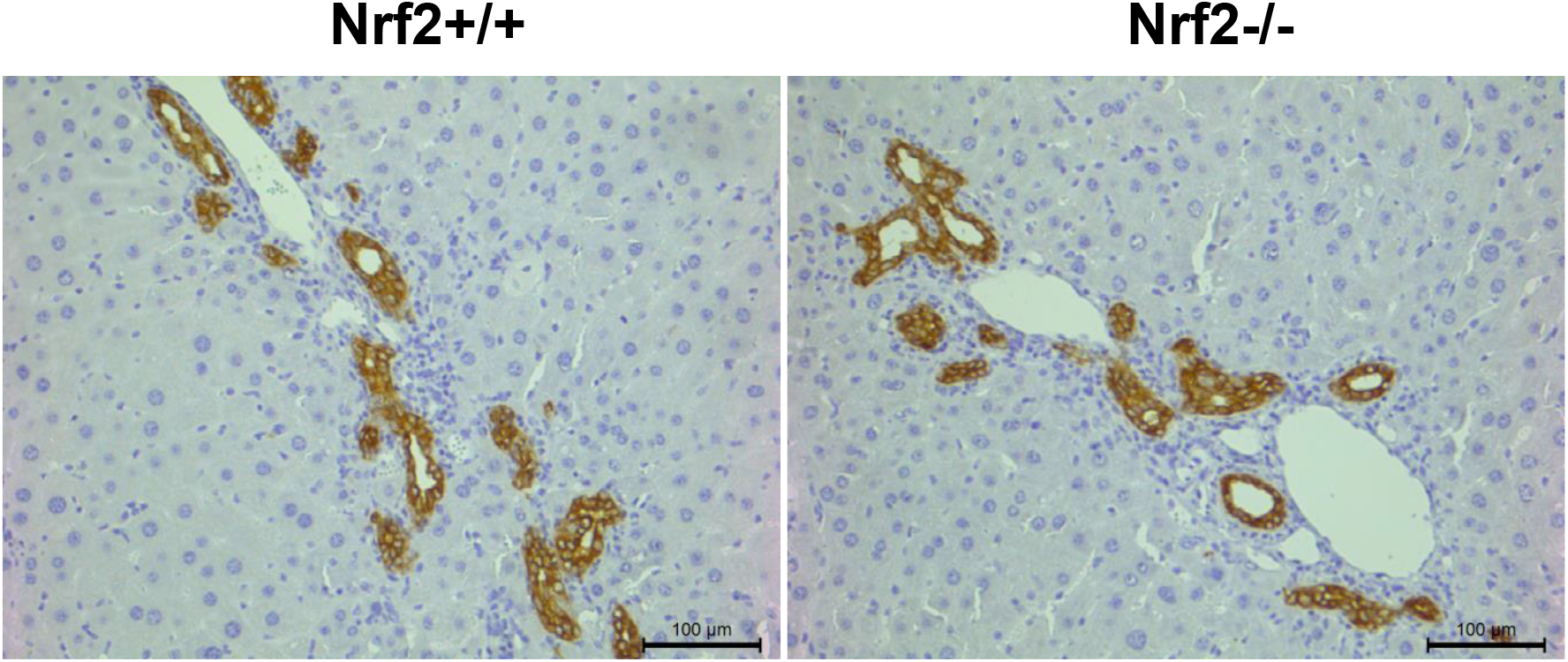
Hepatic distribution of CK19+ cells in Nrf2+/+ and Nrf2-/- mice following bile duct ligation (BDL). Sections were prepared from the livers of mice 40 days after BDL and subjected to CK19 immunostaining. Representative sections show CK19+ cells stained dark brown within the biliary ducts.

**Figure 3.**
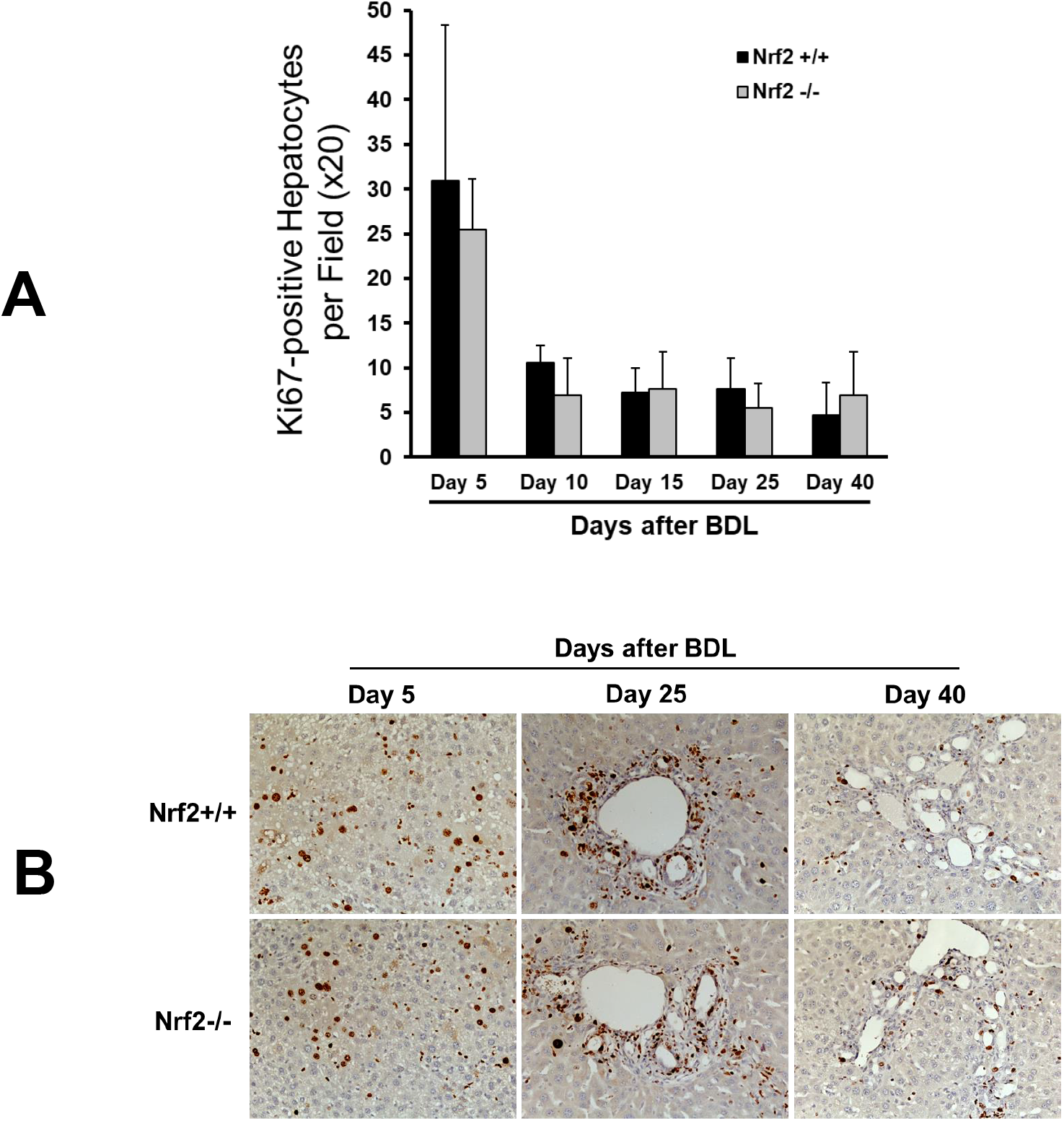
Hepatocyte proliferation induced by bile duct ligation (BDL) in Nrf2+/+ and Nrf2-/- mice. (A) Ki-67 immunostaining was performed using the formalin-fixed and paraffin-embedded liver sections prepared from the samples collected from the livers of mice after BDL or sham operation. Ki67+ hepatocytes were counted at 200x magnification in 5 randomly chosen fields per section. The results are shown as means per field ± standard deviations. \**P* < 0.05 and n = 5. (B) Representative liver sections showing Ki67+ cells with the nuclei stained dark brown.

**Figure 4.**
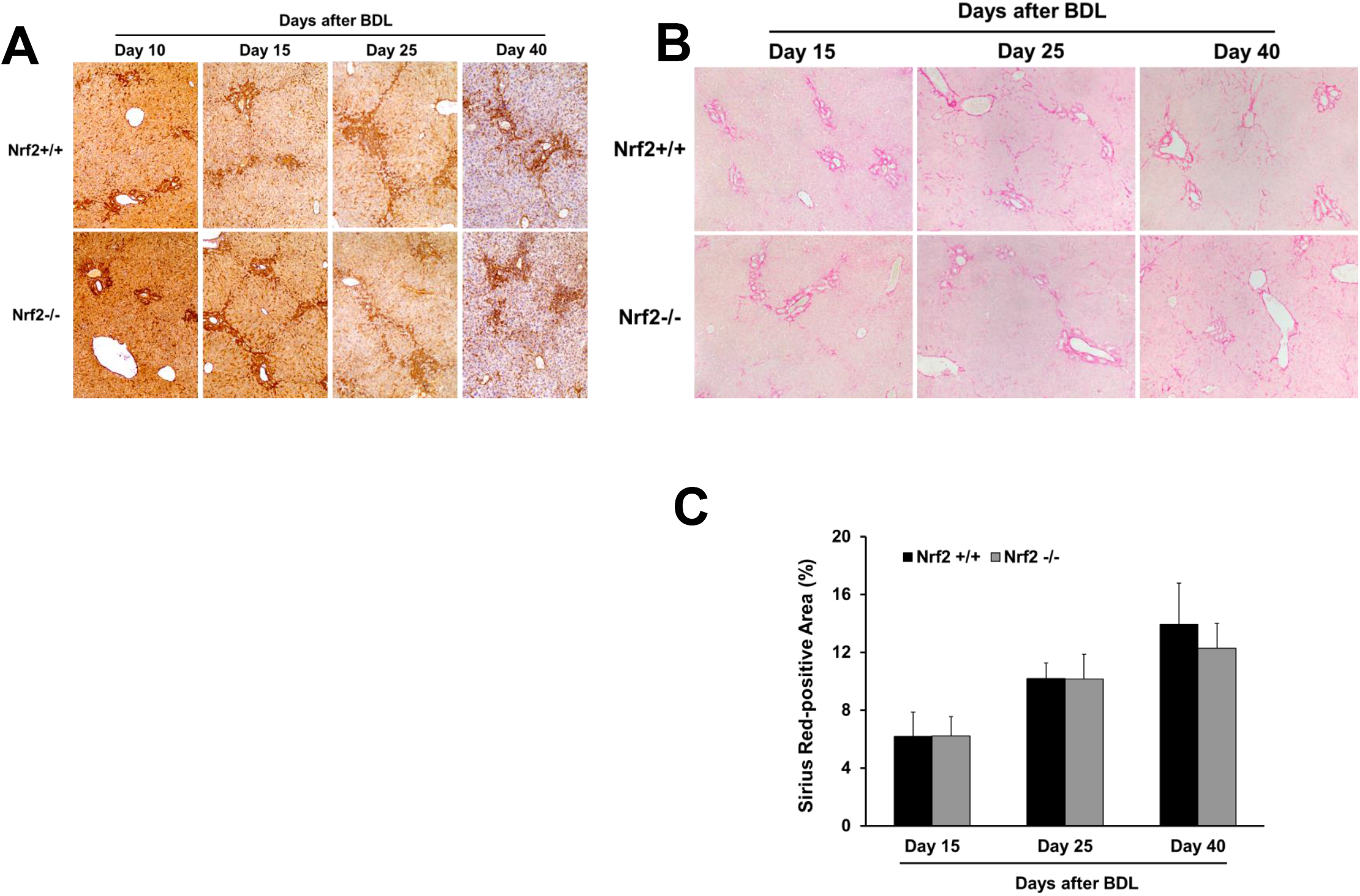
Hepatic fibrotic response to bile duct ligation (BDL) in Nrf2+/+ and Nrf2-/- mice. The liver sections obtained from the livers of mice after BDL or sham operation underwent laminin immunostaining or Sirius Red staining. (1) Representative liver sections with laminin immunostaining. The septal tissue rich in laminin was stained prominently dark brown. (B) Representative liver sections with Sirius Red staining. (C) Sirius Red staining areas were quantified using Image-Pro Plus software (Media Cybernetics, Rockville, MD, USA) at 200x magnification in 5 randomly chosen fields per section. The results are shown as the means of percentages ± standard deviations (n = 5).

### Nrf2 deficiency causes hepatocyte dedifferentiation in response to BDL

We next evaluated the functional state of hepatic Nrf2 by measuring the hepatic expression of NQO1 protein. *Nqo1* is a typical Nrf2 target gene that has been shown to be solely regulated by Nrf2 in BDL-induced cholestasis ^11^. We found that hepatic NQO1 expression was increased as the cholestasis progressed in wild-type mice, which was fully prevented by the absence of Nrf2 (Fig. 5A and 5B). This finding indicated that hepatic Nrf2 exhibited persistent and increased activation in response to BDL, consistent with the result of another study ^11^.

**Figure 5.**
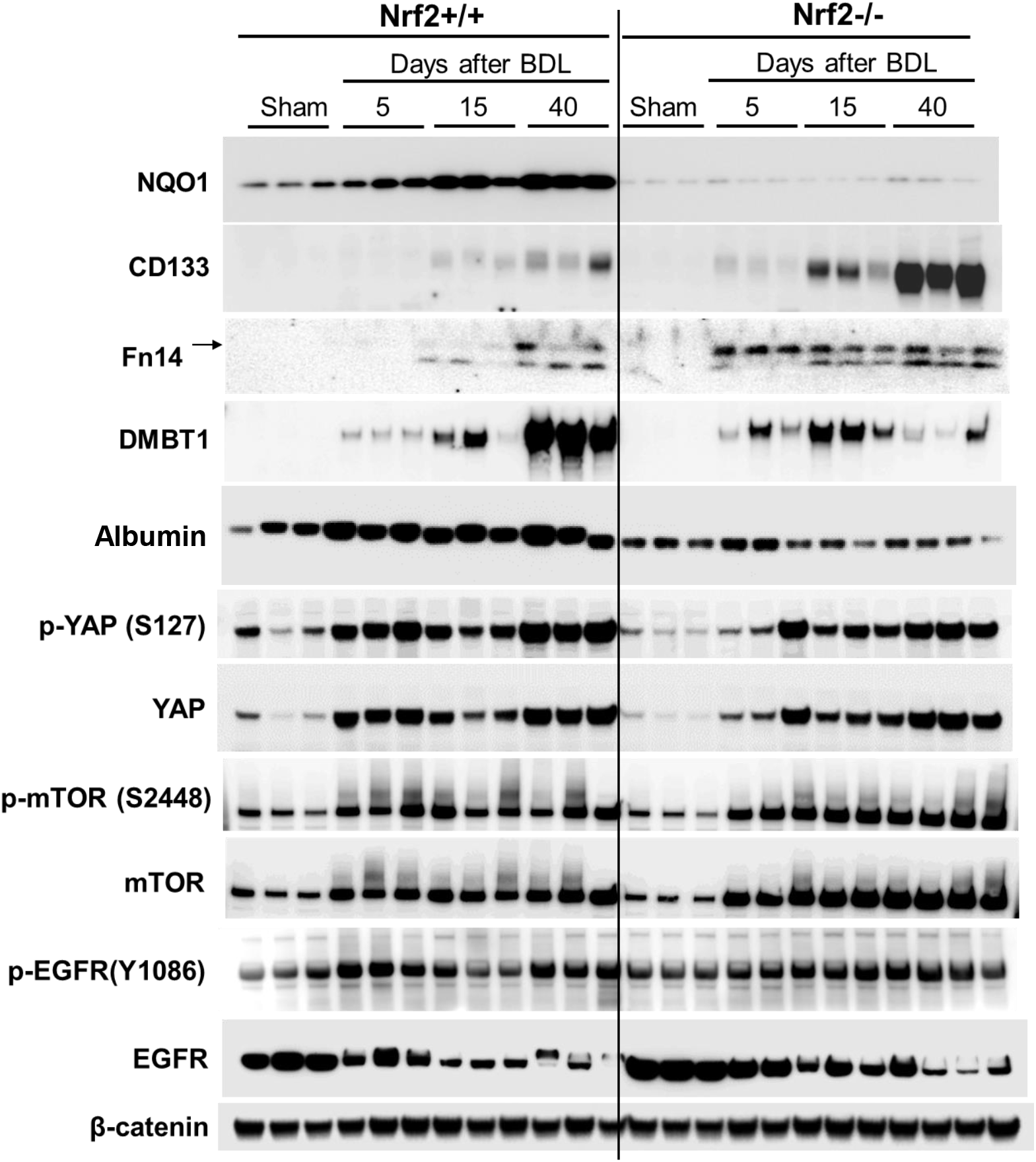
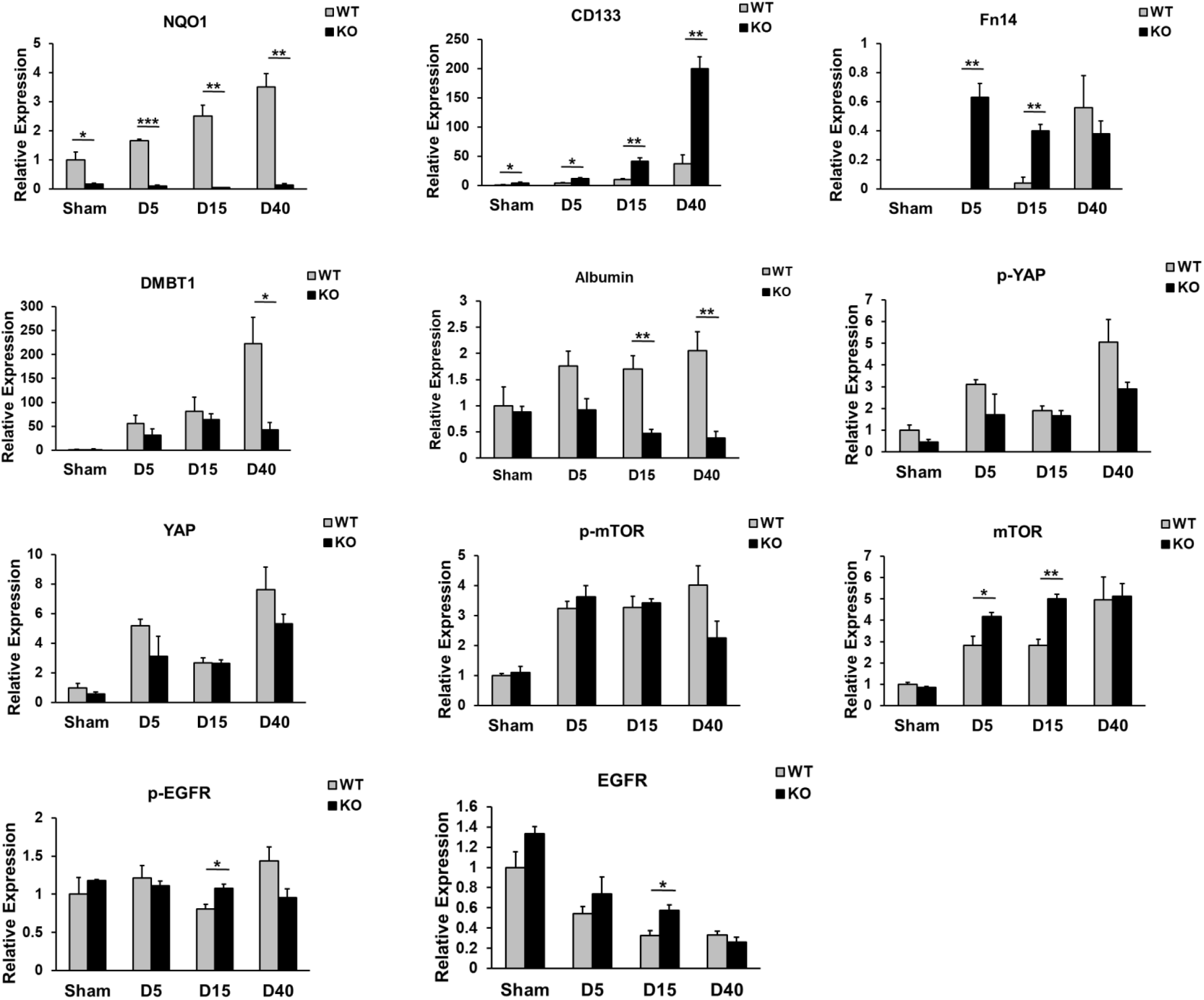
Hepatic expression of a group of proteins in Nrf2+/+ and Nrf2-/- mice after bile duct ligation (BDL). (A) Western blotting was performed with antibodies against the proteins indicated. Liver lysates were prepared from the livers of mice after BDL or sham operation. β-catenon was used as a loading control because it showed the most stable expression over the course of cholestasis relative to regular loading controls screened. Each lane represents a sample from an individual mouse. (B) Densitometry of western blotting. Data are presented as means ± standard deviations normalized with loading controls and relative to the sham controls (day 40 after surgery). \**P* < 0.05, \*\**P* < 0.01, and n = 3.

We next evaluated whether BDL induces a phenotypic response of hepatocytes in an Nrf2-dependent manner by screening a subset of molecules associated with hepatocyte differentiation. As a result, we revealed striking Nrf2-dependent changes in the expression of hepatic progenitor marker CD133 and tumor necrosis factor-like weak induced apoptosis receptor (Fn14), as well as epithelial differentiation factor DMBT1. Western blot (Fig. 5A and 5B) and immunohistochemistry (Fig. 6B and 6C) analyses showed that, as cholestasis progressed to day 15 after BDL, the hepatocytes in the wild-type mice showed very low expression levels of CD133 and Fn14. This observation indicated that hepatocytes underwent slight dedifferentiation. However, during this period, Nrf2 deficiency strengthened the extent of hepatocyte dedifferentiation, as indicated by earlier and stronger induction of CD133 and Fn14 expression. In parallel, after BDL, the hepatocytes induced low DMBT1 expression regardless of Nrf2 expression. Surprisingly, when cholestasis advanced to day 40 post-BDL, the hepatocytes in the wild-type mice not only enhanced Fn14 expression, indicative of increased dedifferentiation, but also drastically upregulated DMBT1 expression. This observation suggested that hepatocytes in the wild-type mice might elevate DMBT1 expression to prevent further dedifferentiation. In contrast, at the same stage of cholestasis, hepatocytes in the Nrf2-null mice retained Fn14 expression but strikingly increased CD133 expression, indicative of more severe dedifferentiation, which was accompanied by the prevention of DMBT1 upregulation. This finding implied that the absence of Nrf2 caused marked hepatocyte dedifferentiation or severe impairment of hepatocyte identity, possibly due to diminished DMBT1 upregulation. Functionally, sham-operated mice produced equivalent amounts of albumin regardless of genotype. However, in response to cholestasis, wild-type hepatocytes persistently increased albumin production, whereas Nrf2-null hepatocytes did oppositely (Fig. 5A-B). By Day 40 following BDL, albumin expression levels in Nrf2-null mice were only about one fourth of that in wild-type mice. On one hand, this was a functional indication of hepatocyte dedifferentiation owing to the lack of Nrf2 during cholestasis. On the other hand, the data showed that Nrf2 deficiency-caused hepatocyte dedifferentiation resulted in adverse consequence in liver function. Collectively, our results demonstrated that Nrf2 modulated hepatocyte phenotypes possibly via the Nrf2/DMBT1 pathway during cholestasis.

**Figure 6.**
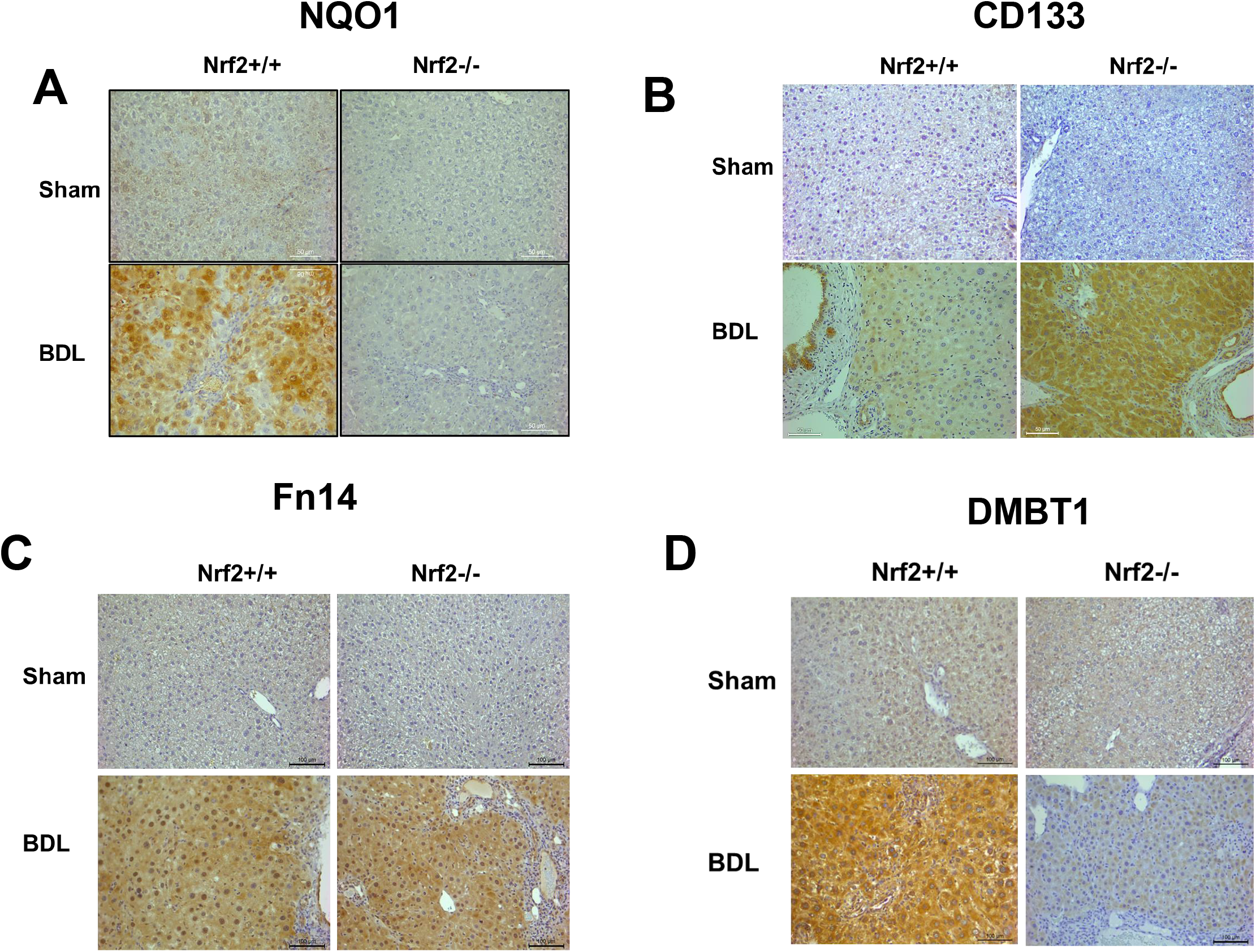
Hepatic distribution of a group of proteins in Nrf2+/+ and Nrf2-/- mice post-bile duct ligation (BDL) or sham operation (Sham). Sections prepared from the livers of mice 40 days after BDL or sham operation were subjected to (A) NQO1, (B) CD133, (C) Fn14, and (D) DMBT1 immunostaining. Representative sections are shown.

### Hepatocytes highly inactivated the yes-associated protein 1 (YAP) and activated the mammalian target of rapamycin (mTOR) signaling to adapt to cholestasis, which is mildly influenced by the lack of Nrf2 expression

YAP has been shown to regulate hepatocyte differentiation ^14,15^. We found that the livers after BDL remarkably and persistently increased the expression of phosphorylated (inactivated) YAP and total YAP compared to those of the sham controls in both genotype groups of mice. However, by day 40 post-BDL, livers of Nrf2-null mice showed a notable reduction in the expression of p-YAP and total YAP compared to those in the wild-type controls, but these differences were not statistically significant (Fig. 5A and 5B). mTOR signaling is central in regulating cellular metabolism and growth ^16^. We observed that the livers in both the wild-type and Nrf2-null mice markedly and continuously upregulated the expression of p-mTOR and total mTOR in response to BDL compared to those of the sham controls. When comparing the two genotype groups of mice, Nrf2 loss of function led to increased total mTOR expression on days 5 and 15 after BDL (Fig. 5A and 5B). Nrf2 is associated with EGFR signaling in the liver ^17^. We found that, compared to the sham controls, BDL did not induce significant changes in hepatic p-EGFR levels but progressively reduced the total EGFR expression in both genotype groups of mice. Taken together, these findings suggested that hepatocytes responded to cholestasis by highly inactivating YAP and activating mTOR signaling and, furthermore, Nrf2 might not play a major role in mediating these responses.

## Discussion

The current study further demonstrates that Nrf2 deficiency does not lead to widely worsened liver damage during BDL-induced cholestasis, in line with the results of other studies. Aleksunes et al. examined the effects of Nrf2 genetic deletion on hepatic NQO1 expression and activity, liver histology, and liver bile acids within 3 days after BDL in mice. They found that Nrf2 knock-out prevented the induction of NQO1 expression, reduced accumulation of liver bile acids, and did not alter the extent of total hepatocellular necrosis ^11^. Weerachayaphorn et al. analyzed serum alanine aminotransferase, liver histology, expression and deposition of hepatic collagen, and expansion and distribution of CK19+ cells 7 days after BDL in wild-type and Nrf2-null mice. They found no significant Nrf2-dependent differences in these assessments ^10^. Here, we evaluated the liver size, biliary ductular reaction, hepatocyte proliferation, and hepatic fibrotic response up to 40 days after BDL and did not observe an overt Nrf2-dependent changes. Taken together, our work and those of others consistently demonstrated that the loss-of-function of Nrf2 does not broadly worsen BDL-induced liver injury. This finding strongly suggests that an unknown mechanism is compensatory to Nrf2 deficiency-caused defects in combating oxidative stress and alleviating liver damage during cholestasis.

The present work also suggests that Nrf2 may not be required for biliary ductular reaction to BDL. Nrf2 has been shown to regulate cholangiocyte differentiation ^18^. Taguchi et al. reported that the forced activation of Nrf2 by liver-specific genetic deletion of both its inhibitor Keap1 and phosphatase and tensin homolog led to abnormal expansion of ductal structures containing cholangiocytes, resulting in severe hepatomegaly, which caused animal death within 1 month after birth. This finding demonstrated that Nrf2 activity needs to be tightly controlled to ensure normal cholangiocyte differentiation during development. BDL induces massive ductular reactions with reprogramming or transdifferentiation of hepatocytes to cholangiocytes in order to adapt to cholestasis ^19^. We observed progressively increased hepatic Nrf2 activity after BDL, suggesting a role for Nrf2 in hepatic adaptive responses to cholestasis. However, we and others did not observe an obvious effect of the genetic deletion of Nrf2 on BDL-induced biliary ductular reaction. This finding implies that Nrf2 plays distinct roles in chalangiocyte differentiation during different physiological and pathological settings.

The most striking finding in our study is that, in response to cholestasis, hepatocytes need to activate Nrf2 to remain in their differentiated state. In this pathological condition, hepatocytes that are deficient in Nrf2 undergo massive dedifferentiation, indicated by a strong CD133+/Fn14+ phenotype in nearly all of them, which is accompanied by striking reduction in albumin production. CD133 is a cholesterol-binding glycoprotein with an unknown function ^20^. Evidence suggests a role for CD133 in the promotion of hepatocellular carcinoma ^21^. It is highly expressed in several types of adult stem/progenitor cells, including hematopoietic, neural, and hepatic progenitors. Thus, it is widely used as a marker for stem/progenitor cells in various tissues ^22^. Several groups have shown that Fn14 is also expressed in hepatic progenitor cells, and its activation stimulates the expansion of these cells ^23-26^. We previously showed that during partial hepatectomy-induced liver regeneration, the absence of Nrf2 caused transient but massive hepatocyte dedifferentiation ^27^. Our previous and current findings strongly support a role for Nrf2 in maintaining hepatocyte phenotypes in diseased livers, which needs to be further investigated.

The present work identified that the Nrf2/DMBT1 pathway is relevant to the maintenance of hepatocyte phenotypes during cholestasis. Evidence supports the notion that when the liver is chronically injured, a subpopulation of hepatocytes exhibit phenotypic plasticity to promote liver repair ^28, 29^. However, most hepatocytes need to stay in the differentiated state to maintain liver function. YAP has been shown to modulate hepatocyte dedifferentiation and differentiation. The activation of YAP expression alone can dedifferentiate hepatocytes to a progenitor phenotype, whereas inactivation of YAP induces differentiation of these cells ^15^. However, we found that BDL led to inactivation of hepatic YAP, independent of Nrf2 and irrespective of the change in hepatocyte phenotype. This observation suggested that YAP may not be a gatekeeper of hepatocyte identity in this experimental setting. Remarkably, we found that, in the late stage of cholestasis, hepatocytes abundantly and Nrf2-dependently expressed DMBT1. It has been demonstrated that DMBT1 is essential for epithelial cell differentiation during development ^30^. Since the *Dmbt1* gene is deleted in several types of epithelial cancers, it is considered a tumor suppressor. Studies have suggested that DMBT1 may play a dominant role in forcing the epithelium to maintain its differentiated state ^31^. Thus, it is highly likely that hepatocytes activate the Nrf2/DMBT1 pathway to adapt to cholestasis in order to prevent hepatocyte dedifferentiation. In fact, when BDL-induced cholestasis progressed to the late stage, hepatocytes in the wild-type mice displayed a tendency of dedifferentiation, which was reflected by weak CD133 and Fn14 expression. However, we assume that the concurrent, high magnitude, and Nrf2-mediated elevation of DMBT1 expression prohibits hepatocytes from dedifferentiation, whereas Nrf2 deficiency blocked DMBT1 upregulation, which resulted in a severe impairment in hepatocyte identity. We are highly interested in elucidating the role and importance of the Nrf2/DMBT1 pathway in modulating hepatocyte phenotypes during injury to livers in our future investigation.

## List of Abbreviations

BDL: bile duct ligation
DMBT1: deleted in malignant brain tumor 1
EGFR: epidermal growth factor receptor
Fn14: fibroblast growth factor-inducible 14
Nrf2: nuclear factor erythroid 2-related factor 2
NQO1: NAD(P)H quinone dehydrogenase 1
YAP: yes-associated protein

---

Graphical Abstract

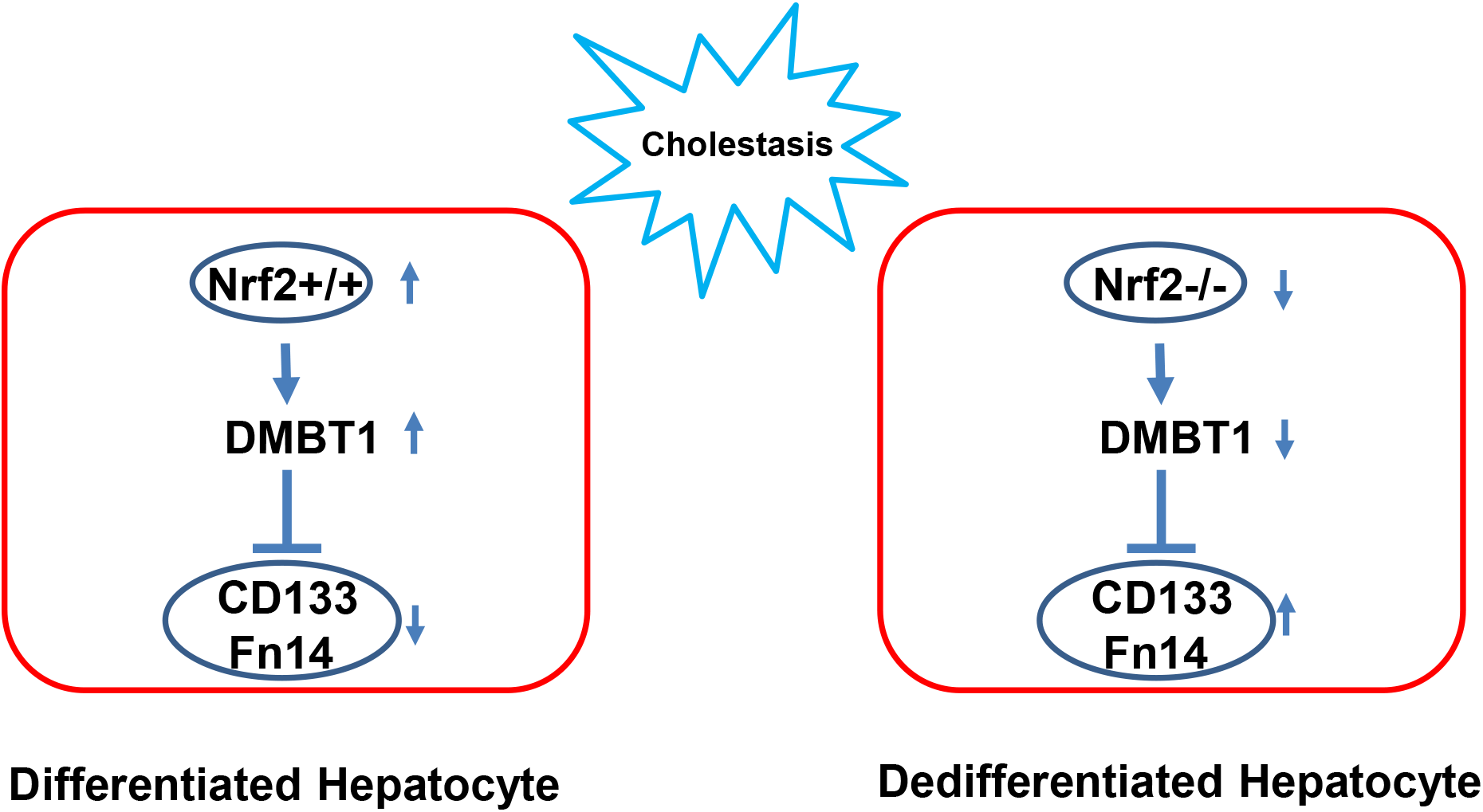

